# *Sodalis praecaptivus* subsp. *spalangiae* subsp. nov., a nascent bacterial endosymbiont isolated from the parasitoid wasp, *Spalangia cameroni*

**DOI:** 10.1101/2024.08.29.610137

**Authors:** Li Szhen Teh, Ian James, Anna Dolgova, Elad Chiel, Colin Dale

## Abstract

An endosymbiotic bacterium of the genus *Sodalis,* designated as strain HZ^T^, was cultured from the parasitoid wasp *Spalangia cameroni*, that develops on pupae of various host flies. The bacterium was detected in *S. cameroni* that developed on house flies, *Musca domestica*, in a poultry facility in Hazon, northern Israel. Following culture, this bacterium displayed no surface motility on LB agar, is rod-shaped and irregular in size, approximating 10-30 nm in diameter and 5-20 μm in length. Phylogenetic analyses revealed that strain HZ^T^ is closely related to *Sodalis praecaptivus* strain HS^T^, a free-living species of the genus *Sodalis* that includes many insect endosymbionts. Although these bacteria maintain >98% sequence identity in shared genes, genomic characterization revealed that strain HZ^T^ has undergone substantial reductive evolution, such that it lacks many gene functions that are maintained in *S. praecaptivus* strain HS^T^. Based on the results of phylogenetic, genomic and chemotaxonomic analyses, we propose that this endosymbiont should be classified in a new subspecies as *Sodalis praecaptivus* subsp. s*palangiae* subsp. nov. The type strain for this new subspecies is HZ^T^ (=ATCC TSD-398^T^=NCIMB 15482^T^). The subspecies *Sodalis praecaptivus* subsp. *praecaptivus* strain HS^T^ is created automatically with the type strain ATCC BAA-2554^T^ (=DSMZ 27494^T^).

## Introduction

In 1999, the genus *Sodalis*, within the family “*Bruguierivoracaceae”*, in the order *Enterobacterales*, in the class *Gammaproteobacteria*, in the phylum *Pseudomonadota*_,_ was established with the isolation, culture, and characterization of *Sodalis glossinidius,* a facultative endosymbiont of the tsetse fly, *Glossina morsitans morsitans* [1]. Since then, symbionts from a wide range of insect taxa have been identified sharing high levels of 16S rRNA gene sequence identity with *S. glossinidius*, but many were not described with validly published species names, due to the absence of laboratory culture, which is often difficult to achieve with fastidious insect endosymbionts [2]. Examples include symbionts of leafhoppers, weevils, mealybugs, stinkbugs, psyllids and lice, in addition to blood-feeding dipteran flies [3–5]. In many cases, the bacteria have been demonstrated to provide beneficial functions for their insect hosts, commonly encompassing the provision of nutrients such as amino acids and vitamins [6–8].

Outside of association with insects, a close relative of *S. glossinidius* was isolated and cultured from a human wound in 2012 and subsequently named *S. praecaptivus* strain HS^T^ [9, 10]. Interestingly, the genomes/gene inventories of many insect-associated *Sodalis*-allied symbionts were found to be subsets of *S. praecaptivus* strain HS^T^, implying that the symbionts are derived from environmental progenitors with genome composition similar to *S. praecaptivus* strain HS^T^, as supported by phylogenetic and comparative genomic analyses [4, 9, 10]. More recently, bacteria isolated from decaying wood samples were also proposed to be members of the genus *Sodalis,* including *S. ligni*, which has been validly published [11], on the basis that they maintain >97% 16S rRNA gene sequence identity with *S. praecaptivus* strain HS^T^ [12]. However, pangenomic comparisons revealed that the wood-decomposing representatives maintain unique functions including lignin degradation and nitrogen fixation that are absent in *S. praecaptivus* strain HS^T^ and insect-associated *Sodalis* spp., which group within a separate clade and maintain genes in common that are not found in deadwood-associated strains [12].

The ability to culture symbionts in the laboratory provides a means to perform experimental infections *in vivo* and to explore molecular interactions in symbiosis in those examples in which bacteria are amenable to genetic modification [2, 13–16]. To this end, *S. praecaptivus* strain HS^T^ and *S. glossinidius* have been used to gain insight into the functions of numerous genes (or networks of genes) including those encoding type III secretion systems, the PhoPQ two-component regulation system that facilitates insect antimicrobial peptide resistance and a quorum sensing system that regulates expression of insect-specific virulence factors [2, 15–17]. Furthermore, in a recent study, *S. praecaptivus* strain HS^T^ was engineered to replicate and replace the function of a native (long-established) symbiont of a grain weevil, yielding a novel, synthetic symbiosis replete with mutualistic functionality and vertical symbiont transmission [18]. Implementation of these techniques also fuels opportunities to utilize symbiotic bacteria to control insect vector-borne diseases, by engineering symbionts to interfere with processes of pathogen colonization and/or transmission by insects [19].

In this study, we focus on the cultivation of a novel member of the *Sodalis* genus that was recently identified as a symbiont of a parasitoid wasp, *Spalangia cameroni* (Hymenoptera: Spalangiidae) collected in Hazon, Israel [20]. Subsequently, it was determined to have high fidelity of maternal transmission in *S. cameroni* and to persist in a range of host tissues [21]. Phylogenetic analyses revealed an extremely close relationship between this symbiont and *S. praecaptivus* strain HS^T^, suggesting a very recent origin of the symbiotic association. Based on this observation, we reasoned that the symbiont of *S. cameroni* should be amenable to laboratory culture, given that it likely retains a large quotient of gene functions found in the free-living relative, *S. praecaptivus* strain HS^T^. Thus, the objective of this work was to perform cultivation, characterization, phylogenetic and genomic analysis of strain HZ^T^, establishing a new, experimentally tractable system to explore interactions between insects and symbiotic bacteria.

## Methods

### House fly rearing

Adult house flies were held in net cages with water and a diet of sugar, milk powder and egg yolk powder mixture (2:2:1 by weight, respectively). The larvae were reared on a medium of wheat bran mixed with calves’ food pellets, wetted with water to 60-65% moisture. The flies were maintained at 26 ±1 °C, 60 ±20% relative humidity (RH), and 14:10 light/dark photoperiod.

### Parasitoid rearing

*Spalangia cameroni* was collected in 2014 from an egg-laying poultry facility in Hazon, Israel (32°54’25.8“N; 35°23’49.0”E), using sentinel pupae of *Musca domestica* as previously described [22]. The parasitoids were subsequently reared on house fly pupae, under conditions of 25 ±1 °C, 60 ±20% RH, and 14:10 light/dark photoperiod.

### Strain HZ^T^ cultivation

Twenty *Spalangia cameroni* adults that harbor strain HZ^T^ were chilled on ice and placed in 1% bleach (NaOCl) for 3 min, on ice, to sterilize their external integument. All the subsequent steps were carried out in a sterile flow hood. The wasps were rinsed three times with sterile double deionized water (DDW), placed in groups of five in 0.5 ml sterile tubes containing 100 µl LB medium with polymyxin B sulfate (25 μg/ml) and 0.5% L-cysteine (as an antioxidant), and were homogenized using a sterile pestle. Each lysate was added to a 15 ml sterile tube containing 10 ml of sterile molten LB medium with 0.1% low-melting point agarose and 25 μg/ml polymyxin B sulfate at 37 °C and mixed by inversion to distribute the bacteria (the polymyxin B sulfate was added after the medium was sterilized and cooled down below 60 °C).

Each mix was poured into a sterile glass tube with a solid plug at the bottom of 1% agar containing 5 mg/ml thioglycolate and sealed with its sterile cap. The tubes were maintained vertically at 27 °C without disturbing once set. The incubated tubes were checked daily until a band of cells was visible about 1 cm below the surface of the medium at a reduced oxygen tension. One microliter of agar and cells extracted from this region was checked by PCR using *Sodalis*-specific primers targeting a *Sodalis*-specific *marR* homolog (locus tag Sant_4061). An additional 5 μl was seeded on LB agar plates containing 25 μg/ml polymyxin B sulfate, 0.16 mM IPTG and 0.032 mg/ml X-Gal, and incubated at 27 °C. These additives were included to reduce contamination and facilitate identification based on the observation that *S. praecaptivus* strain HS^T^ maintains an intact *lacZ* homolog and (along with *S. glossinidius*) is known to be resistant to polymyxin B (mediating antimicrobial peptide resistance in insects [17]). The plates were checked daily and blue colonies that grew were tested by colony PCR for the presence of the *marR* (Sant_4061) gene. The primers used were: F5’-GCGTATCTATCACCGCCTGA-3’, R5’-CATTGCAGACTTCGCTGAAC-3’, generating a 359 bp amplicon. PCR conditions comprised an initial denaturation cycle of 2 min at 95 °C, followed by 30 cycles of 30 sec at 94 °C, 30 sec at 60 °C, 30 sec at 72 °C, and a final extension step of 5 min at 72 °C. The PCR product was separated by electrophoresis on a 1.5% agarose gel, stained with RedSafe (iNtRON biotechnology) and visualized by UV transillumination. The PCR products were purified using a PCR purification kit (Qiagen) and Sanger-sequenced to verify their identity. Each PCR reaction included a positive control (a lysate of *Sodalis*-infected *Spalangia cameroni*) and a negative control (PCR mix with no added lysate). The positive control lysates were prepared as follows: a single wasp was homogenized in 40 μl of buffer (10 mM Tris-Cl pH 8.2, 1 mM EDTA, 25mM NaCl) containing 2 mg/ml proteinase K (Invitrogen). The lysates were incubated for 15 min at 65 °C, then 10 min at 95 °C and kept at −20 °C until further use. Strain HZ^T^ has been deposited in public culture collections at the American Type Culture Collection (TSD-398^T^) and the National Collection of Industrial, Food and Marine Bacteria (NCIMB 15482^T^).

### Glycerol stock preparation

Each blue colony that formed on the agar plates prior to testing by PCR, was streaked on a new LB agar plate containing polymyxin B sulfate, IPTG and X-Gal (concentrations specified above) and incubated under the same conditions as described previously. Single colonies that grew on these plates were used to inoculate 5 ml liquid LB + 25 μg/ml polymyxin B in 15 ml sterile tubes and incubated vertically without shaking until bacterial growth was visible. The 5 ml cultures were transferred to 50 ml sterile tubes and 10 ml fresh media were added. Another 25 ml fresh media were added after one week, and after 7-10 additional days. For cryopreservation, 80% sterile glycerol was added to the bacterial cultures to provide a final concentration of 20%. The glycerol stocks were aliquoted into Eppendorf tubes (1 ml each) and stored at −80 °C.

### Growth of strain HZ^T^ at different temperatures

Strain HZ^T^ from a glycerol stock was streaked on an LB agar plate containing polymyxin B sulfate, IPTG and X-Gal (concentrations as above) and incubated at 27°C. From the colonies that grew on this plate, a single colony was picked to inoculate a 10 ml starter culture, incubated at 27 °C, without shaking. When the OD_600nm_ of this starter culture had reached 0.14, 0.5 ml aliquots were transferred to 12 tubes, each containing 49.5 ml liquid LB +25 μg/ml polymyxin B. Groups of three tubes (experimental replicates) were incubated at different temperatures: 23 °C, 28 °C, 33°C and 37 °C, without shaking. Growth of strain HZ^T^ was monitored by OD_600nm_ (turbidity) measurements taken every 48 h in the first four days, and then every 24 h between days 5-14. The turbidity measurements were made using 1 ml cuvettes (polystyrene 10×4×45 mm, SARSTEDT) with the NanoDrop One spectrophotometer (Thermo Fisher Scientific).

### Microscopy

Liquid cultures of *Sodalis praecaptivus* strain HS^T^ and strain HZ^T^ were grown in LB + 25 μg/ml polymyxin B at 30 °C until OD_600nm_ reached 0.1. For fluorescence microscopy, 1.5 ml of culture was pelleted at 8,000 ×g for 10 min. Cells were then resuspended in 100 μl of phosphate-buffered saline (PBS) and stained by addition of 1 μl of 100 μg/ml FM4-64 (Molecular Probes) and visualized on an Echo Revolve microscope using a 100× fluorite objective and TRITC filter cube. For transmission electron microscopy, *S. praecaptivus* strain HS^T^ and strain HZ^T^ cultures were grown in LB + 25 μg/ml polymyxin B at 30 °C until OD_600nm_ reached 0.03 and 15 ml of each culture was pelleted at 3,000 ×g for 10 min. Bacteria were resuspended in 200 µl of freshly prepared fixative (32% paraformaldehyde in Live Cell Imaging solution; Invitrogen) and incubated for 2 h at room temperature. Cells were pelleted again before resuspension in 50 μl water and 3 μl of cell suspension was then deposited onto a glow discharged formvar-carbon-coated grid. The cells adsorbed on the grid were passed over three consecutive drops of 0.1% phosphotungstic acid, pH 7.0 and excessive fluid on the grid was removed by blotting with filter paper. Air-dried specimens were then examined with a transmission electron microscope (JOEL JEM-1400Plus) operated at 120 kV at the University of Utah Electron Microscopy Core Laboratory. Transmission electron micrographs were analyzed using Gatan Digital Micrograph software version 3.0.

### PCR amplification of 16S rRNA gene sequence

The 16S rRNA gene sequence of strain HZ^T^ was amplified by PCR using primers 27F (F5’-AGAGTTTGATCMTGGCTCAG-3’) and 1492R (F5’-GGTTACCTTGTTACGACTT-3’), using a 2X PCR Master Mix (Thermo Scientific) and cycling conditions consisting of a 5 min initial denaturation at 94 °C, 30 cycles of denaturation (1 min at 94 °C), annealing (1 min at 50 °C) and extension (1 min at 72 °C), followed by a final extension at 72 °C for 5 min. The resulting amplicon was purified using AxyPrep Mag PCR Clean-Up beads (Axygen) and sequenced using the Sanger method.

### Genome sequencing

A 40 ml liquid culture of the strain HZ^T^ was grown in LB + 25 μg/ml polymyxin B at 30 °C until OD_600nm_ reached 0.1. Cells were pelleted at 9,000 ×g for 10 min and transferred into 180 μl Buffer ATL. DNA was extracted using a Qiagen Blood and Tissue Kit protocol (Qiagen) following the manufacturer’s protocol for Gram-negative bacteria. The genomic DNA was treated with 1 μl RNaseA for 15 min at room temperature and purified using Ampure XP purification beads following the manufacturer’s protocol (Axygen). Purified DNA was eluted in 50 μl nuclease-free water, the DNA concentration was determined using a NanoDrop Lite Spectrophotometer (Thermo Scientific) and DNA quality was assessed by electrophoresis.

Library construction and sequencing was performed using the Illumina Nextera DNA Flex Library Preparation Kit and Illumina NovaSeq 6000 S4 reagent Kit v1.5 to generate 150-base paired-end reads in accordance with the manufacturer’s protocols. Sequencing was performed on an Illumina NovaSeq 6000 instrument at the University of Utah Huntsman Cancer Institute High-Throughput Genomic Core Facility, yielding 14.8M raw paired-end reads. Reads were trimmed using BBDuk [23] at both ends with minimum quality of Q30 and any reads <100-bases were discarded. The resulting 5.2M trimmed paired-end reads were assembled using Tadpole [23] with default parameters.

For Nanopore sequencing, genomic DNA was isolated from 50 ml of a strain HZ^T^ culture grown without shaking in LB media at 30 °C with 25 µg/ml polymyxin B sulfate to an OD_600nm_ of 0.09. High molecular weight (HMW) genomic DNA was isolated using a Circulomics Nanobind CBB Big DNA Isolation Kit (Circulomics) following the “Nanobind HMW DNA Extraction Gram-Negative Bacteria” protocol. The resulting soluble and insoluble fractions were collected and sequenced separately by preparing libraries using the SQK-LSK110 library preparation kit (Oxford Nanopore Technologies) from 1.3 µg of each DNA sample. Resulting libraries were loaded onto FLO-MIN106 flow cells and sequenced in the Mk1C sequencing device (Oxford Nanopore Technologies) for 20 h (soluble fraction) and 22 h (insoluble fraction). Primary acquisition of data and real-time basecalling was carried out using the graphical user interface MinKNOW v2.0 (Oxford Nanopore Technologies, https://community.nanoporetech.com), using default parameters, and reads passing the quality filter were extracted after sequencing for downstream analysis.

Raw Nanopore FAST5 reads were basecalled using MinKNOW Core module v4.4.3 (https://community.nanoporetech.com) employing High Accuracy Calling mode, and resulting fastq reads from each sequencing run were filtered to remove reads < 60-kbp in size. The Nanopore reads from each run were then combined and assembled *de novo* using Flye (version 2.8.3) [24] with the -nano raw argument. The resulting contigs were subjected to polishing using Medaka v1.5.0 (https://github.com/nanoporetech/medaka), using default parameters, and four iterations of Pilon (v1.24) [25]. In each iteration of Pilon (v1.24) [25], MiniMap2 (v2.23) [26] was used to map Illumina reads to the assembly, using the preset for short single-ended reads without splicing, and the option to generate output in a SAM format, following which SAMtools (v1.12) [27] was used to sort and index the alignment created by MiniMap2 (v2.23) [26], using default parameters for indexing, and finally Pilon (v1.24) [25], was used to polish the assembly based on the aforementioned alignment. Genome annotation was performed on 21 contigs derived from the assembly of Nanopore reads from strain HZ^T^ using the automated NCBI Prokaryotic Genome Annotation Pipeline (PGAP) version 6.7 [28]. A custom script was generated to identify pseudogenes, tRNAs, rRNAs, phage and transposon genes via keyword search in the annotation file to facilitate data visualization.

### Comparative genomics

Illumina reads from strain HZ^T^ were aligned to the *S. praecaptivus* strain HS^T^ chromosome and plasmid sequences (GenBank CP006569 and CP006570) using Geneious Prime 2023.1.2 with default parameters to facilitate comparative analysis of the gene inventories. Average nucleotide identity (ANI) and coverage (shared genomic content) between strain HZ^T^ contigs and the *S. praecaptivus* strain HS^T^ chromosome and plasmid was estimated using the OrthoANIu algorithm [29]. Digital DNA:DNA hybridization (dDDH) was performed using the same query sequences with the genome-to-genome distance calculator [30], using the recommended formula (#2).

### Phylogenetic Analyses

Whole genome phylogenetic analysis was performed using the Type Strain Genome Server (TYGS; https://tygs.dsmz.de/) [31] following importation of requisite whole genome sequences from GenBank, including strain HZ^T^, other *Sodalis* spp. and *Pectobacterium carotovorum* (as outgroup). For 16S rRNA gene sequence and ribosomal protein-coding gene-based analyses, all strain HZ^T^ sequences were extracted from a consensus sequence derived from the alignment of Illumina reads to the *S. praecaptivus* strain HS^T^ genome sequence. Orthologous sequences from other bacteria, encompassing numerous *Sodalis*-allied symbionts and related environmental isolates, were obtained from NCBI. All sequence alignments for phylogenetic analyses were generated using MUSCLE v5.1 [32] in Geneious Prime and inspected manually. For the alignment of concatenated ribosomal protein-coding genes, a translation alignment (using MUSCLE v5.1 [32]) was generated from 47 genes comprising *rpsJ*, *rplCDWB*, *rpsS*, *rplV*, *rpsC*, *rplP*, *rpmC*, *rpsQ*, *rplNXE*, *rpsH*, *rplFR*, *rpsE*, *rplO*, *rpsMKD*, *rplQ*, *rplY*, *rplT*, *rpmI*, *rplS*, *rpsP*, *rpmA*, *rplU*, *rplI*, *rpsR*, *rpsF*, *rpsI*, *rplM*, *rplLJAK*, *rpmF*, *rpmGB*, *rpsLG*, *rpsB*, *rpsO*, and *rpsU*. Maximum likelihood (ML) phylogenies were then inferred using RAxML v8.2.11 [33], using a GTR+GAMMA substitution model with 10,000 bootstrap replicates. For both the 16S rRNA gene sequence and ribosomal protein-coding gene trees, sequences from *Pectobacterium carotovorum* were designated as the outgroup for each rooted tree.

### Phenotypic and Metabolic Features

API 50 CH test strips (bioMérieux) were utilized to examine carbon source utilization by strain HZ^T^ and strain HS^T^ (for comparative purposes). Following the manufacturer’s instructions, bacteria were grown in 40 ml LB + 25 μg/ml polymyxin B at 30 °C until OD_600nm_ reached 0.12, pelleted by centrifugation (6,000 × *g*., 10 min) and resuspended to generate suspensions with turbidities of 2 McFarland units for testing. Following inoculation into API 50 CHL medium, the strip wells were covered with mineral oil and incubated at 30 °C. API strips were inspected daily and the results were documented after 48 h.

## Results and Discussion

### Strain HZ^T^ cultivation

As reported in a subsequent section of these results, whole genome sequencing revealed that the genome of strain HZ^T^ includes intact homologs of the machinery comprising the *lac* operon, and maintains the necessary genetic machinery to grow in the presence of antimicrobial peptides such as polymyxin B. Hence, we exploited these features in our procedure to cultivate strain HZ^T^ from *Spalangia cameroni*.

Strain HZ^T^ was isolated from members of a laboratory population of *S. cameroni* that have maintained this symbiont for several years [20]. Bacterial growth was visible 5-10 days post-inoculation (Fig. 1A) and was enhanced on plates incubated under microaerophilic conditions. Two results were indicative of the success of isolating and culturing the bacterial symbiont from its host. First, the PCR with *Sodalis marR*-specific primers yielded a single abundant amplicon with a sequence sharing 99.7% identity with the *Sodalis praecaptivus* strain HS^T^ ortholog (GenBank accession CP006569). Second, blue colonies on IPTG/X-Gal agar plates were formed (Fig. 1B) and colony PCR assays provided the same results as detailed above, indicating that strain HZ^T^ also produces β-galactosidase.

**Fig. 1.**
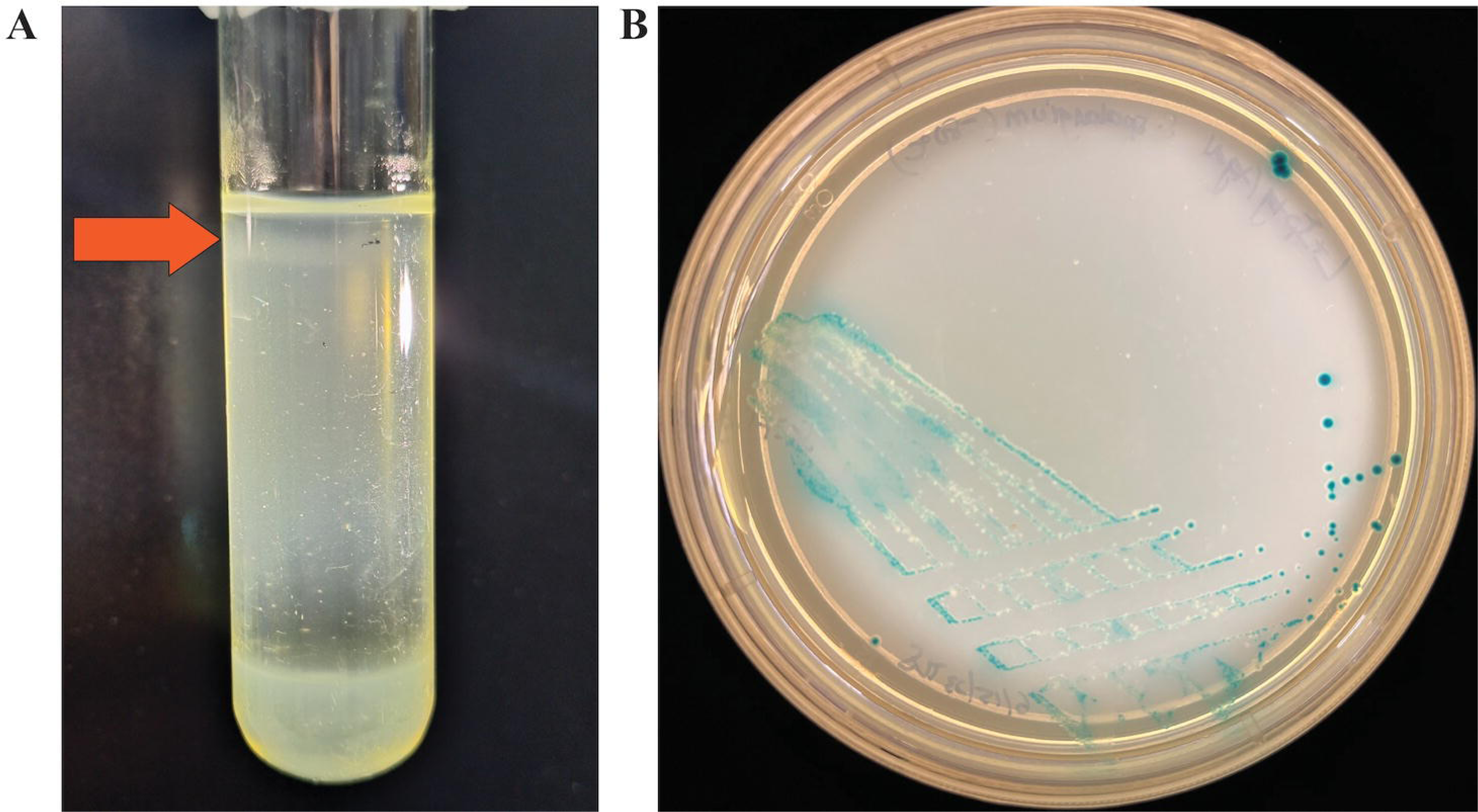
**A.** Strain HZ^T^ growing in the microaerobic zone of a semi-solid LB agar redox gradient, highlighted with orange arrow. **B.** Strain HZ^T^ growing on LB agar plates with IPTG, X-Gal and polymyxin B sulfate.

### Growth of Strain HZ^T^ at different temperatures

Overall, the highest growth rate was observed at 28°C with a maximal OD_600nm_ of 0.18, followed by 23°C and 33°C (Fig. S1). Growth at 37°C was substantially reduced relative to the other three temperatures and only achieved an OD_600nm_ of 0.03. In the first four days, we observed steady growth at all four temperatures, which was optimal at 28°C. Between days 4-14, the growth rates curtailed and remained relatively stationary, possibly due to the activities of endogenous phages as explained later.

### Microscopy

Colonies of strain HZ^T^ were variable in size, white in the absence of X-Gal, circular and glistening with entire edges. They displayed no surface motility on LB agar. Fluorescence imaging revealed that the cells are filamentous, relative to *S. praecaptivus* strain HS^T^, approximating 10-30 nm in diameter and 5-20 μm in length (Fig. 2). Transmission electron microscopy confirmed these findings and revealed the presence of lateral flagella on both *S. praecaptivus* strain HS^T^ and strain HZ^T^, although these structures were less abundant on the latter (Fig. 2). Further, phage particles were observed in cultures of strain HZ^T^, having a 54 nm head and long, contractile tail (Fig. S2).

**Fig. 2.**
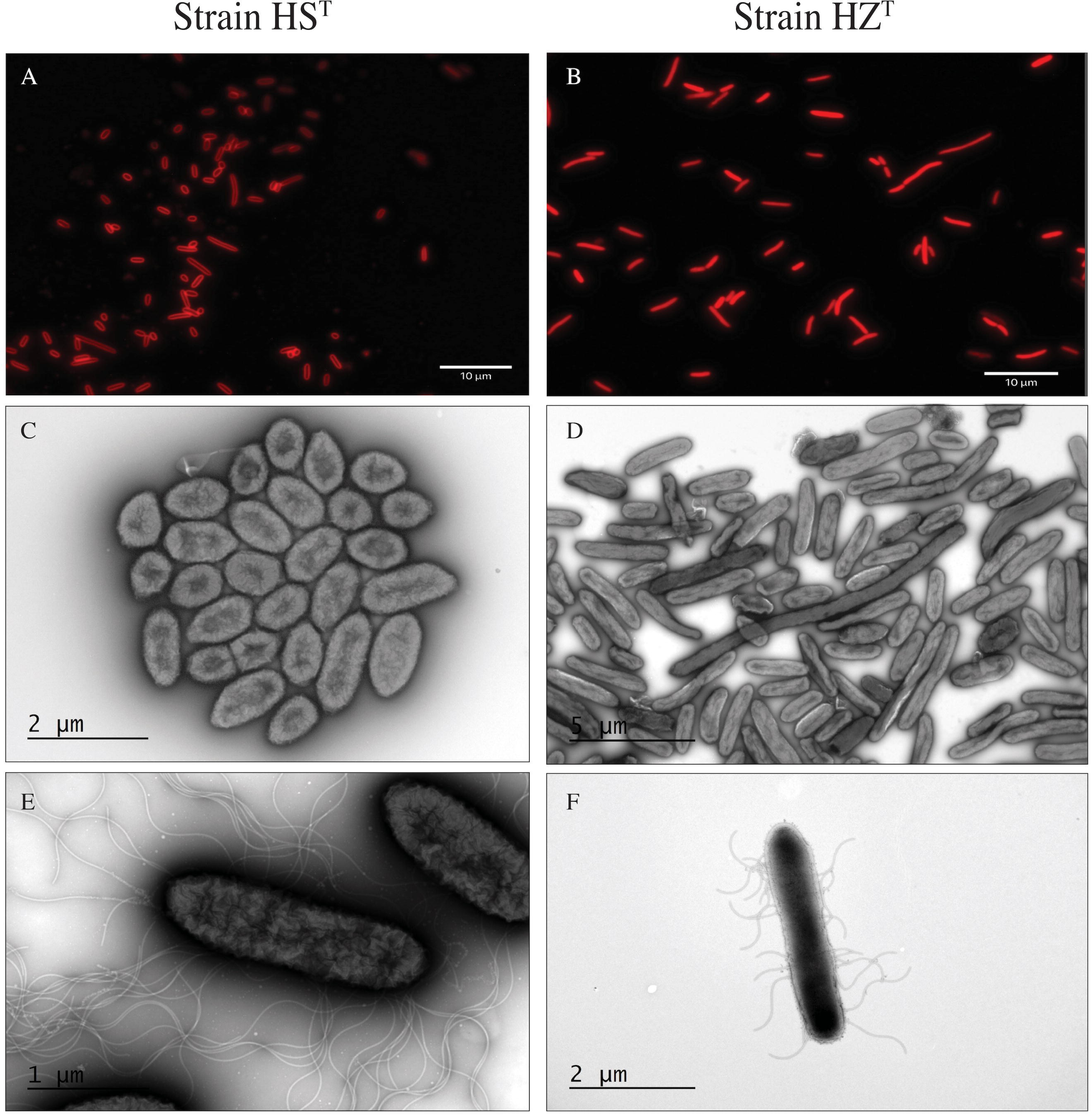
**A.** Micrographs of cultured *S. praecaptivus* strain HS^T^ taken using fluorescence microscopy following staining with FM4-64. **B.** Micrographs of cultured strain HZ^T^ taken using fluorescence microscopy following staining with FM4-64. **C and E.** Transmission electron micrographs of *S. praecaptivus* strain HS^T^ cells at low and high magnifications, respectively (see scale bars for reference). **D and F**. Transmission electron micrographs of strain HZ^T^ cells at low and high magnifications, respectively (see scale bars for reference).

### Genome sequencing and analysis

The strain HZ^T^ Illumina sequence reads assembled using Tadpole yielded 1,393 contigs with N50 size of 8,522 bp, indicating that the assembly was highly fragmented. Inspection of the sequences at the contig ends using BLAST analyses revealed that the contigs were terminated by repetitive sequences sharing high levels of identity with insertion sequences and phages. Proliferation of mobile genetic elements is known to occur frequently in the early stages following establishment of symbiotic associations [34] and is documented for other *Sodalis*-allied symbionts [35, 36]. Since high levels of repetitive DNA diminish the utility of short-read assembly, we also performed Nanopore sequencing, followed by polishing/error correction using the Illumina reads. The resulting sequence assembly (GenBank GCA_039646275.1; Table S1) yielded a single circular chromosome of 5.24 Mb with 56.8% G+C content and 408-fold mean coverage. An additional 20 contigs were obtained comprising putative extrachromosomal elements ranging in size from 0.52-kbp to 231-kbp and coverage ranging from 3-fold to 513-fold. Due to the large numbers of mobile DNA elements in the strain HZ^T^ genome, these 20 contigs likely represent a combination of assembly artefacts in addition to canonical (replicative) plasmids, DNA from phage particles located in bacterial cells, and possibly other non-replicating elements arising from mobile element activity. Preliminary annotation of the strain HZ^T^ genome was conducted using the PGAP (version 6.7) [28] and deposited under accession JBDHPY000000000 and assembly number GCA_039646275.1. Further manual annotation will be required to definitively characterize and annotate extrachromosomal elements and the extensive array of mobile DNA insertions in this genome. In order to determine if the relationship between strain HZ^T^ and *S. praecaptivus* strain HS^T^ merits placement of strain HZ^T^ in the same species, we computed average nucleotide identity (ANI) and performed DNA:DNA hybridization (dDDH) between the two strains. These yielded estimates of ANI= 98.5% and dDDH=85.4% that would typically be considered high enough to merit placement of the bacteria in the same species. However, comparison of gene content between *S. praecaptivus* strain HS^T^ and strain HZ^T^ reveals key distinctions (Table S1) and shows that strain HZ^T^ maintains only a subset of the gene inventory of *S. praecaptivus* strain HS^T^, evidenced by (1) an average genome coverage (determined by orthoANIu) of only 50.2% and (2) absence of alignment of strain HZ^T^Illumina reads to numerous regions of the *S. praecaptivus* strain HS^T^ genome (Fig. S3), noting that the same reads align contiguously to the strain HZ^T^ Nanopore assembly (Fig. S4). Furthermore, strain HZ^T^ harbors substantial numbers of novel, mobile DNA elements and predicted pseudogenes (Table S1; Fig. S5). Based on this insight it appears that, like many other insect symbionts including *Sodalis* spp., strain HZ^T^ has undergone reductive genome evolution, involving deletions, gene inactivating mutations and accumulation/proliferation of mobile genetic elements [4, 5, 8, 35–37]. Thus, while intact coding sequences shared between *S. praecaptivus* strain HS^T^ and strain HZ^T^ maintain high sequence identity, indicative of recent common ancestry, *S. praecaptivus* strain HS^T^ and strain HZ^T^ may differ substantially in terms of gene inventories and resulting phenotypic properties. Therefore, we propose that strain HZ^T^ is classified in a new subspecies as *Sodalis praecaptivus* subsp. *spalangiae* subsp. nov. This is justified on the basis that it maintains >98% sequence identity in gene sequences shared with *S. praecaptivus* strain HS^T^, yet harbors many pseudogenes and mobile DNA elements, in addition to occupying a distinct ecological niche.

### Phylogenetic Analyses

Phylogenomic analysis was performed with the Type Strain Genome Server (TYGS) [31] using reference genome sequences derived from *Sodalis*-allied symbionts, including strain HZ^T^, several free-living *Sodalis* spp. and an outgroup, *Pectobacterium carotovorum* (Fig. 3). The resulting tree shows that strain HZ^T^ is the closest known, insect-associated relative of *S. praecaptivus* strain HS^T^ discovered to date again supporting the notion that the association between the wasp *Spalangia cameroni* and strain HZ^T^ is very recent in origin. Additional phylogenetic analyses (Fig. S6-8) utilizing the PCR-derived 16S rRNA gene sequence (deposited in NCBI GenBank under accession number OR464179.1) and 47 ribosomal protein-coding genes (deposited in NCBI GenBank under accession numbers PP273189-235), yield similar results. Notably, the 16S rRNA gene sequence tree shows that the strain HS^T^ sequence shares higher sequence identity with strain HZ^T^ than with the sequence derived from the symbiont of the weevil *Archarius roelofsi*, which was formerly the closest known insect-associated relative of strain HS^T^ [9]. All three trees also provide strong support for a clade comprising *S. praecaptivus* strain HS^T^ and *Sodalis*-allied insect symbionts excluding *Sodalis* spp. isolated from deadwood and soil, as reported in another study [12]. In addition, the 16S rRNA gene sequence derived from the cultured strain HZ^T^ shares 100% sequence identity with the 16S rRNA gene sequence derived from the wasp host reported previously [20] (GenBank KY985293), indicating that the cultured isolate matches the *Sodalis* symbiont reported in that study.

**Fig. 3.**
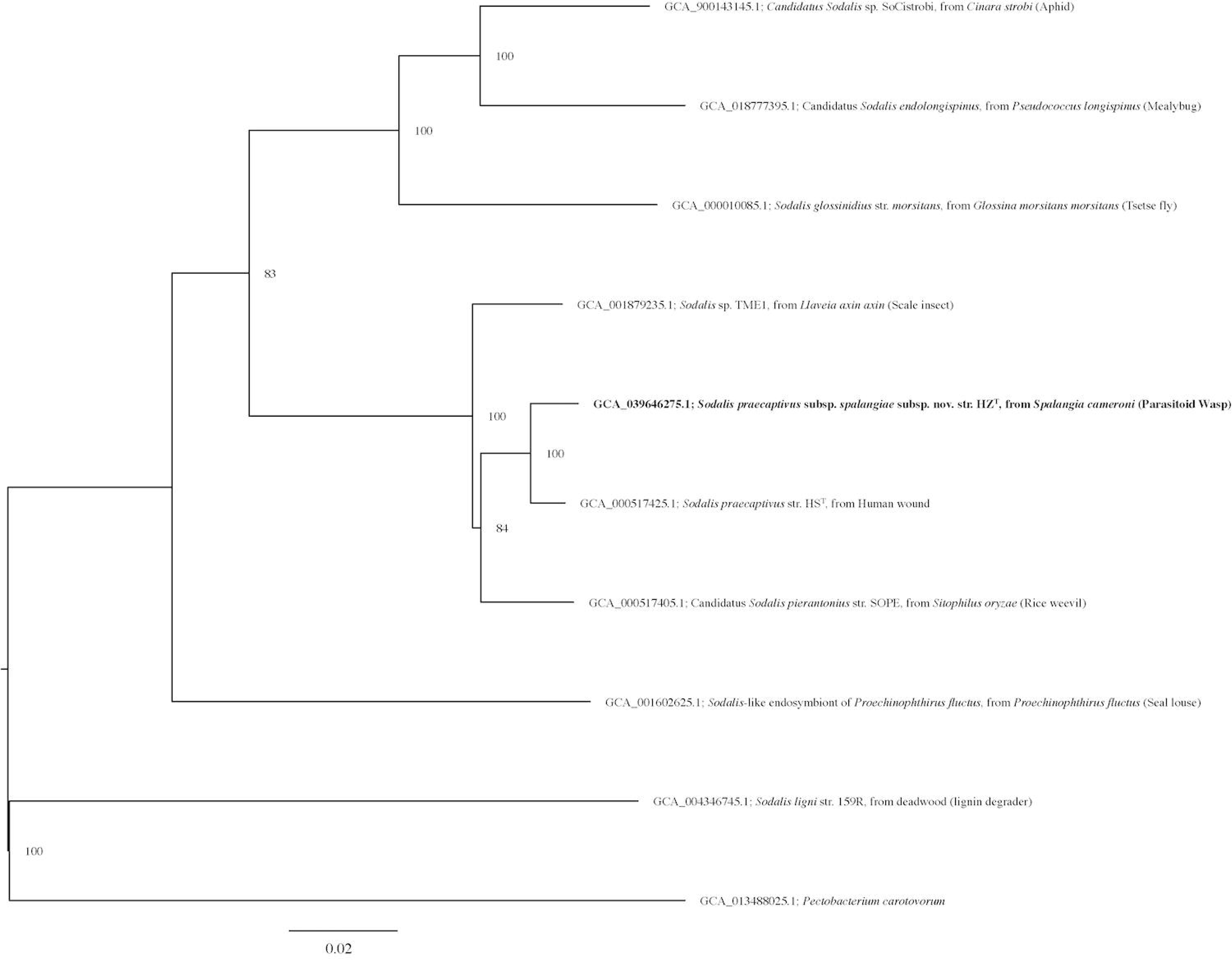
Phylogenomic tree based on genome sequences of strain HZ^T^ and related bacteria in the TYGS (https://tygs.dsmz.de/). The midpoint rooted tree was inferred with FastME 2.1.6.1 [31] from GBDP distances calculated from genome sequences. The branch lengths are scaled in terms of GBDP distance formula d_5_. The numbers above branches are GBDP pseudo-bootstrap support values >60□% from 100 replications, with an average branch support of 95.3□%.

### Phenotypic and Metabolic Features

API 50 CH strip tests revealed the carbon source utilization capabilities and preferences of *S. praecaptivus* strain HS^T^ and strain HZ^T^ (Table S2). The results derived for *S. praecaptivus* strain HS^T^ were in agreement with those obtained from a previous study [10]. The phenotypic test results also suggest that strain HZ^T^ maintains a subset of the metabolic properties found in strain HS^T^, again consistent with the notion that strain HZ^T^ has undergone genomic and functional degeneration following the transition to symbiotic life. This includes loss of the ability to utilize six of the 23 carbon sources utilized by *S. praecaptivus* strain HS^T^ in the API 50 CH testing regime (L-arabinose, D-ribose, D-xylose, L-xylose, D-lactose, and D-melibiose). Notably, strain HZ^T^ was found to lack the ability to utilize lactose in spite of the fact that it produces blue colonies on plates with IPTG and X-Gal. This may be indicative of loss of function of some other gene(s) required for lactose utilization or it may be due to a change in the specificity of the ß-galactosidase enzyme. Also, observation of the API strips at time intervals following inoculation revealed that strain HZ^T^ favors the utilization of N-Acetylglucosamine over other carbon sources (including glucose) present in the test, as evidenced by it having the most rapid color change.

Interestingly, this mirrors findings documented for carbon source utilization by *S. glossinidius* [1] which also showed a metabolic preference for N-Acetylglucosamine, noted to be a constituent of chitin, the abundant exoskeletal component of insects. In addition, we found that strain HZ^T^, unlike strain HS^T^, does not grow on M9 minimal medium (with either glucose or N-Acetylglucosamine provided as a carbon source), suggesting that it is auxotrophic for one or more amino acids, vitamins and/or co-factors.

### Proposal of Novel Subspecies within *S. praecaptivus*

Levels of nucleotide sequence identity, based on discrete alignments and estimation of ANI and dDDH, between strain HZ^T^ and *S. praecaptivus* strain HS^T^, are above the thresholds typically used to define species boundaries [38, 39], yet strain HZ^T^ lacks genes and phenotypic properties that are maintained in the closely related *S*. *praecaptivus* strain HS^T^, as evidenced by comparative genomics and phenotypic testing. Therefore, we propose that strain HZ^T^ be designated as a new subspecies of *Sodalis praecaptivus* as *Sodalis praecaptivus* subsp. *spalangiae* subsp. nov. By virtue of this change, we also propose that strain HS^T^ be designated *Sodalis praecaptivus* subsp. *praecaptivus* subsp. nov.

### Emended Description of *Sodalis praecaptivus*

*Sodalis praecaptivus* (prae.cap.ti′vus, L. prep. *prae* before; L. masc. adj. *captivus* taken prisoner; N.L. masc. adj. *praecaptivus* not yet taken prisoner).

Cells are aerobic, Gram-stain-negative, non-spore-forming short rods with 2-10 mm in length and 0.5–1.5 mm in diameter. Colonies are white and circular when grown at 30 °C on LB agar plates. In the API 50 CH assay, the following carbon sources are utilized at 30 °C: glycerol, D-arabinose, D-galactose, D-glucose, D-fructose, D-mannose, L-rhamnose, D-mannitol, D-sorbitol, N-acetylglucosamine, D-trehalose, xylitol, D-lyxose, D-tagatose, L-fucose, potassium gluconate and potassium-5-ketogluconate. In contrast, the following carbon sources are not utilized at 30 °C: erythritol, D-adonitol, methyl-ß-D-xylopyranoside, L-sorbose, dulcitol, methyl-_a_-D-mannopyranoside, arbutin, salicin, D-maltose, L-arabitol and potassium 2-ketogluconate. Produce ß-galactosidase (forming blue colonies with the presence of X-Gal) and are polymyxin B resistant.

The chromosomal sizes are in the range of 4.71-5.25 Mb and the G+C content is 56.8-57.5%. The type strain is HS^T^ (=DSM 27494^T^=ATCC BAA-2554^T^), isolated from a human patient in the USA. The GenBank/EMBL/DDBJ accession numbers for the genome sequence of strain HS^T^ are CP006569.1 (chromosome) and CP006570.1 (plasmid), those for the 16S rRNA gene and *groEL* gene sequences are JX444565 and JX444566, respectively.

### Description of Sodalis praecaptivus subsp. praecaptivus subsp. nov

*Sodalis praecaptivus* subsp. *praecaptivus* subsp. nov. (prae.cap.ti′vus, L. prep. *prae* before; L. masc. adj. *captivus* taken prisoner; N.L. masc. adj. *praecaptivus* not yet taken prisoner).

The description of this subspecies follows the species description of *Sodalis praecaptivus* with the following additions.

Colonies are mucoid and grown to 1–2 mm in diameter after 24 h of growth at 30 °C on LB agar plates. Growth is observed on LB agar, MacConkey agar, Columbia blood agar, MM blood agar and M9 minimal salt agar from 25 °C to 37 °C. With shaking (200 rpm), liquid cultures reach an OD_600nm_ of ∼0.4-0.5 after 16 h in LB media at 30°C. Exhibits swarming motility on LB agar and grow in LB broth at an optimal pH of 5.5 and a maximum NaCl concentration of 3.5 %. In the API 50 CH assay, L-arabinose, D-ribose, D-xylose, L-xylose, esculin ferric citrate, D-cellobiose, D-lactose, D-melibiose, D-melezitose, D-raffinose, D-turanose, D-fucose, and D-arabitol are utilized as carbon sources at both temperatures, 30 °C and 35 °C. Starch and glycogen are utilized at 30 °C, while inositol, methyl _a_-D-glucopyranoside, amygdalin, D-saccharose (sucrose), inulin and gentiobiose are utilized at 35 °C only. Produces catalase and chitinase.

Strain adheres to the species description. This subspecies type strain is the same as the species type strain HS^T^ (=DSM 27494^T^=ATCC BAA-2554^T^), isolated from a human patient in the USA. The GenBank/EMBL/DDBJ accession numbers for the complete genome sequence of strain HS^T^ are CP006569.1 (chromosome) and CP006570.1 (plasmid), those for the 16S rRNA gene and *groEL* gene sequences are JX444565 and JX444566, respectively.

### Description of *Sodalis praecaptivus* subsp. *spalangiae* subsp. nov

*Sodalis praecaptivus* subsp. *spalangiae* subsp. nov. (spa.lan’gi.ae. N.L. gen. n. *spalangiae*, of the wasp genus *Spalangia*).

The description of this subspecies follows the species description of *Sodalis praecaptivus*. However, this subspecies can be distinguished from *Sodalis praecaptivus* subsp. *praecaptivus* subsp. nov. based on different growth properties. Cells are non-motile in both liquid LB medium and on the surface of LB agar. Grows microaerobically in semi-solid agar but tolerates atmospheric oxygen sufficiently to form irregularly sized colonies after 5-7 days of incubation on LB agar at 30 °C. Cells grow slowly in the absence of shaking in LB broth reaching an OD_600nm_ of ∼0.1 after four days. Colonies are glistening with entire edges. Capable of growth at 23-33 °C with optimum at 28 °C. In the API 50 CH assay, this subspecies cannot utilize L-arabinose, D-ribose, D-xylose, L-xylose, D-lactose, and D-melibiose as carbon sources at 30 °C. Demonstrates faster growth on N-Acetylglucosamine relative to glucose.

The type strain is HZ^T^ (NCIMB 15482^T^=ATCC TSH-398^T^) and was isolated from the parasitoid wasp, *Spalangia cameroni* (Hymenoptera: Spalangiidae), collected in an egg-laying poultry facility in Hazon, Israel in April-May 2014, where the wasps develop on pupae of the housefly, *Musca domestica* (Diptera: Muscidae). The reads described in this study have been deposited in the NCBI sequence read archive under the accession numbers SRR25556807 (Illumina) and SRR28852625 (Nanopore). The 16S rRNA gene sequence and ribosomal protein-coding gene sequences used for phylogenetic analyses are deposited in NCBI GenBank under the accession numbers OR464179.1 and PP273189-235). The genome sequence is deposited in GenBank under the accession number JBDHPY000000000 and assembly number GCA_039646275.1

## Abbreviations

LB: Luria-Broth
ML: maximum likelihood
IS: insertion sequence
ANI: average nucleotide identity
dDDH: digital DNA:DNA hybridization
PGAP: Prokaryotic Genome Annotation Pipeline
OD_600nm_: optical density at 600nm wavelength
DNA: deoxyribonucleic acid
PCR: polymerase chain reaction
ATCC: American Type Culture Collection
NCIMB: National Collection of Industrial Food and Marine Bacteria
DSMZ: Leibniz Institute DSMZ-German Collection of Microorganisms and Cell Cultures GmbH
rRNA: ribosomal ribonucleic acid
DDW: double deionized water
IPTG: isopropyl β-D-1-thiogalactopyranoside
X-Gal: 5-bromo-4-chloro-3-indolyl-β-D-galactopyranoside
Tris-Cl: tris hydrochloride
EDTA: ethylenediaminetetraacetic acid
PBS: phosphate-buffered saline
TYGS: Type Strain Genome Server
tRNA: transfer ribonucleic acid
NCBI: National Center for Biotechnology Information
MM: Mitsuhashi and Maramorosch medium
G+C: Guanine + Cytosine
RH: relative humidity
FM4-64: *N*-(3-triethylammoniumpropyl)-4-(6-(4-(diethylamino)phenyl)hexatrienyl)pyridinium dibromide
TRITC: tetramethylrhodamine isothiocyanate
Q30: Phred quality score 30
HMW: high molecular weight
BLAST: Basic Local Alignment Search Tool.

## Acknowledgements

We gratefully acknowledge Willisa Liou and the University of Utah Electron Microscopy Facility for the electron microscopy images. We thank University of Utah Huntsman Cancer Institute High-Throughput Genomics Core Facility for assistance with high throughput sequencing and Diane Downhour in the Department of the Human Genetics at the University of Utah School of Medicine, for assistance with Nanopore sequencing.

## Author contributions

L.S.T.: conceptualization, investigation, analysis, visualization, writing – original, review and editing

S.R.S.: conceptualization, investigation, analysis, writing-original, review and editing

I.J.: conceptualization, investigation, analysis, visualization, writing-original, review and editing

A.D.: investigation, analysis, writing – review

E.C.: conceptualization, supervision, project administration, analysis, writing-original, review and editing

C.D.: conceptualization, supervision, project administration, analysis, visualization, writing-original, review and editing

