## Supplemental Table 1-2, Supplemental Figure 1-6 for "*Sodalis praecaptivus* subsp. *spalangiae* subsp. nov., a nascent bacterial endosymbiont isolated from the parasitoid wasp, *Spalangia cameroni*": Supplemental updated aug27 v2.pdf

Li Szhen Teh[1,#], Sarit Rohkin Shalom[2,#], Ian James[1], Anna Dolgova[2], Elad Chiel[2], Colin Dale[1]

[1] School of Biological Sciences, University of Utah, Salt Lake City, UT 84112, USA
[2] Department of Biology and Environment, University of Haifa-Oranim, Tivon 36006, Israel
[#] These authors contributed equally to this publication.

**Keywords:** endosymbiont, insect, culture, degenerative evolution, fastidious, insertion sequences, nascent symbiosis

**Repositories:** The Illumina reads generated by the sequencing efforts described in this study have been deposited in the NCBI sequence read archive under the accession number SRR25556807. The 16S rRNA sequence is deposited in NCBI GenBank under the accession number OR464179.1. The ribosomal protein-coding gene sequences are deposited in NCBI GenBank under the accession numbers PP273189 through PP273235. The Nanopore-derived assembly of the genome has been deposited in NCBI Genbank under accession number JBDHPY000000000 and assembly number GCA_039646275.1

**Contents:**

| | | |
|---|---|---|
| Supplementary Table 1 | Genome features of strain HZ$^T$ and *Sodalis praecaptivus* strain HS$^T$ | 2 |
| Supplementary Table 2 | Carbon source utilization for strain HZ$^T$ and *Sodalis praecaptivus* strain HS$^T$ | 3 |
| Supplementary Figure 1 | Strain HZ$^T$ growth at different temperatures | 4 |
| Supplementary Figure 2 | Micrograph of phage particle from strain HZ$^T$ | 5 |
| Supplementary Figure 3 | Histogram of coverage of strain HZ$^T$ reads aligned to *S. praecaptivus* strain HS$^T$ | 6 |
| Supplementary Figure 4 | Histogram of coverage of strain HZ$^T$ reads aligned to strain HZ$^T$ | 7 |
| Supplementary Figure 5 | Strain HZ$^T$ gene categorization | 8 |
| Supplementary Figure 6 | Phylogeny based on rRNA genes | 9 |
| Supplementary Figure 7 | Phylogeny based on ribosomal protein-coding genes | 10 |
| Supplementary Figure 8 | Patristic distance matrix from ribosomal protein-coding gene phylogeny | 11 |

**Table S1.** Composition of the Nanopore assembly of the genome of strain HZ$^T$ and comparison of key features with *S. praecaptivus* strain HS$^T$.

| | *Sodalis praecaptivus* strain HS$^T$ | strain HZ$^T$ |
|---|---|---|
| Total genome size | 5,159,425 bp | 6,134,628 bp |
| Chromosome size | 4,709,528 bp | 5,246,808 bp |
| Extrachromosomal elements | 1 | 20 |
| Extrachromosomal element(s) total size | 449,897 bp | 887,820 bp |
| Extrachromosomal element 1 size | 449,897 bp | 231,212 bp |
| Extrachromosomal element 2 size | | 103,666 bp |
| Extrachromosomal element 3 size | | 101,152 bp |
| Extrachromosomal element 4 size | | 72,231 bp |
| Extrachromosomal element 5 size | | 60,293 bp |
| Extrachromosomal element 6 size | | 53,777 bp |
| Extrachromosomal element 7 size | | 47,917 bp |
| Extrachromosomal element 8 size | | 44,344 bp |
| Extrachromosomal element 9 size | | 41,717 bp |
| Extrachromosomal element 10 size | | 32,170 bp |
| Extrachromosomal element 11 size | | 29,416 bp |
| Extrachromosomal element 12 size | | 21,090 bp |
| Extrachromosomal element 13 size | | 15,287 bp |
| Extrachromosomal element 14 size | | 13,007 bp |
| Extrachromosomal element 15 size | | 6,363 bp |
| Extrachromosomal element 16 size | | 5,082 bp |
| Extrachromosomal element 17 size | | 3,705 bp |
| Extrachromosomal element 18 size | | 3,649 bp |
| Extrachromosomal element 19 size | | 1,222 bp |
| Extrachromosomal element 20 size | | 520 bp |
| CDSs (total) | 4,357 | 6,138 |
| No. of protein-coding genes | 4,331 | 5,881 |
| No. of rRNAs (5S, 16S, 23S) operons | 8, 7, 7 | 8, 7, 7 |
| No. of tRNAs | 76 | 70 |
| No. of pseudogenes | 26 | 257 |
| Chromosomal G+C content (mol%) | 57.5 | 56.8 |
| Overall G+C content (mol%) | 57.1 | 56.1 |

**Table S2**. Comparison of carbon source fermentation between *Sodalis praecaptivus* strain $HS^T$ and strain $HZ^T$ using API 50 CH. +, Positive; w, Weakly positive; -, Negative

| Carbon source | *Sodalis praecaptivus* strain $HS^T$ | strain $HZ^T$ |
|---|---|---|
| Glycerol | w | w |
| Erythritol | - | - |
| D-Arabinose | + | + |
| L-Arabinose | + | - |
| D-Ribose | + | - |
| D-Xylose | + | - |
| L-Xylose | w | - |
| D-Adonitol | - | - |
| Methyl-β-D-Xylopyranoside | - | - |
| D-Galactose | + | + |
| D-Glucose | + | + |
| D-Fructose | + | + |
| D-Mannose | + | + |
| L-Sorbose | - | - |
| L-Rhamnose | + | w |
| Dulcitol | - | - |
| Inositol | - | - |
| D-Mannitol | + | + |
| D-Sorbitol | + | + |
| Methyl-α-D-Mannopyranoside | - | - |
| Methyl-α-D-Glucopyranoside | - | - |
| N-AcetylGlucosamine | + | + |
| Amygdalin | - | - |
| Arbutin | - | - |
| Esculin ferric citrate | - | - |
| Salicin | - | - |
| D-Cellobiose | - | - |
| D-Maltose | - | - |
| D-Lactose (bovine origin) | + | - |
| D-Melibiose | + | - |
| D-Saccharose (sucrose) | - | - |
| D-Trehalose | + | + |
| Inulin | - | - |
| D-Melezitose | - | - |
| D-Raffinose | - | - |
| Amidon (starch) | - | - |
| Glycogen | - | - |
| Xylitol | + | + |
| Gentiobiose | - | - |

| Carbon source | *Sodalis praecaptivus* strain HS$^T$ | strain HZ$^T$ |
|---|---|---|
| D-Turanose | - | - |
| D-Lyxose | + | + |
| D-Tagatose | + | w |
| D-Fucose | - | - |
| L-Fucose | w | w |
| D-Arabitol | - | - |
| L-Arabitol | - | - |
| Potassium Gluconate | + | w |
| Potassium 2-Ketogluconate | - | - |
| Potassium 5-Ketogluconate | + | w |

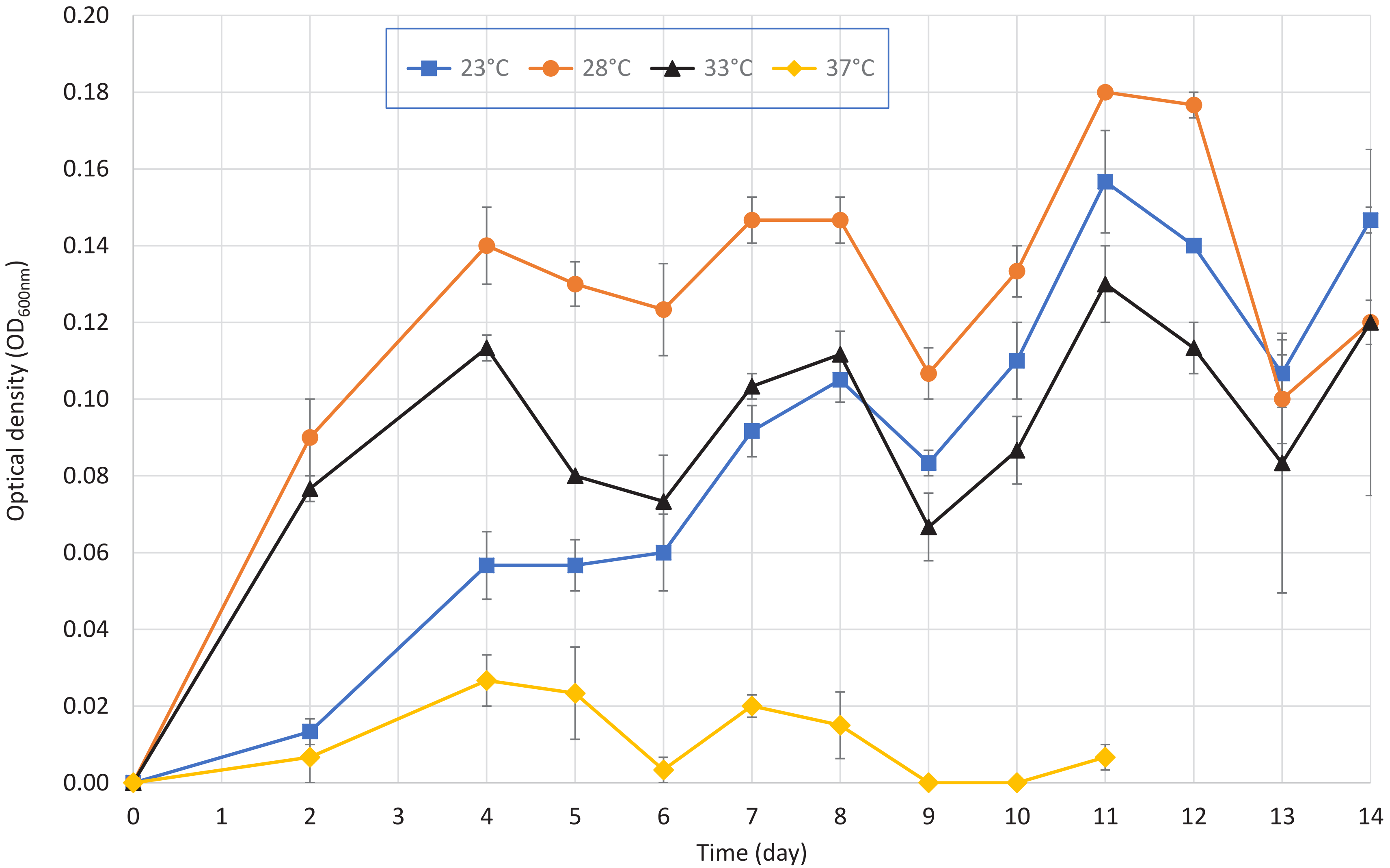


**Fig. S1.** Growth curves of strain $HZ^T$ in liquid LB medium with polymyxin B sulfate at different temperatures. Each data point represents a mean derived from three experimental replicates with error bars representing standard deviations.

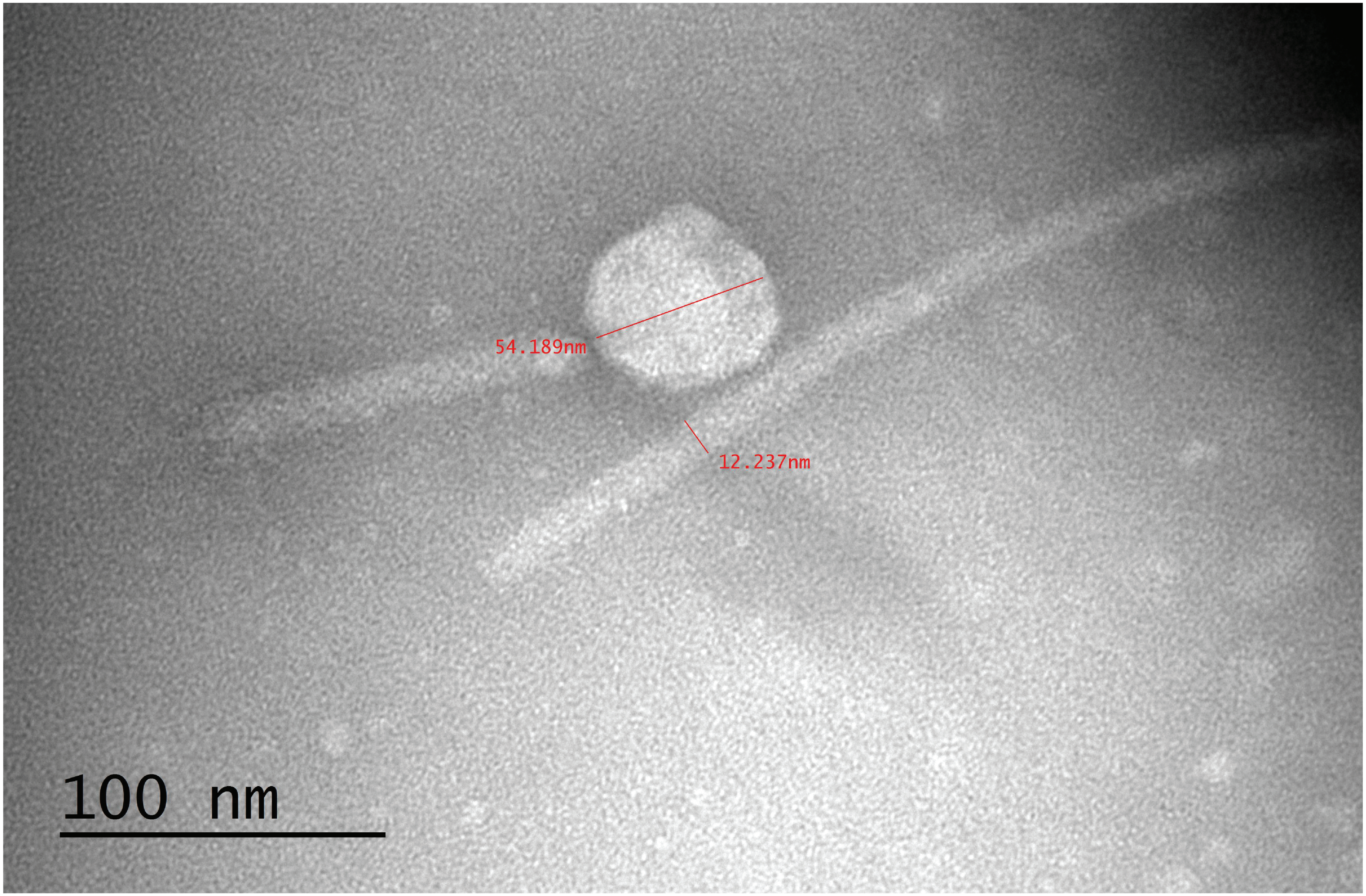


**Fig. S2.** Transmission electron micrograph of phage particle observed in the culture of strain $HZ^T$ that was utilized in Fig. 2.

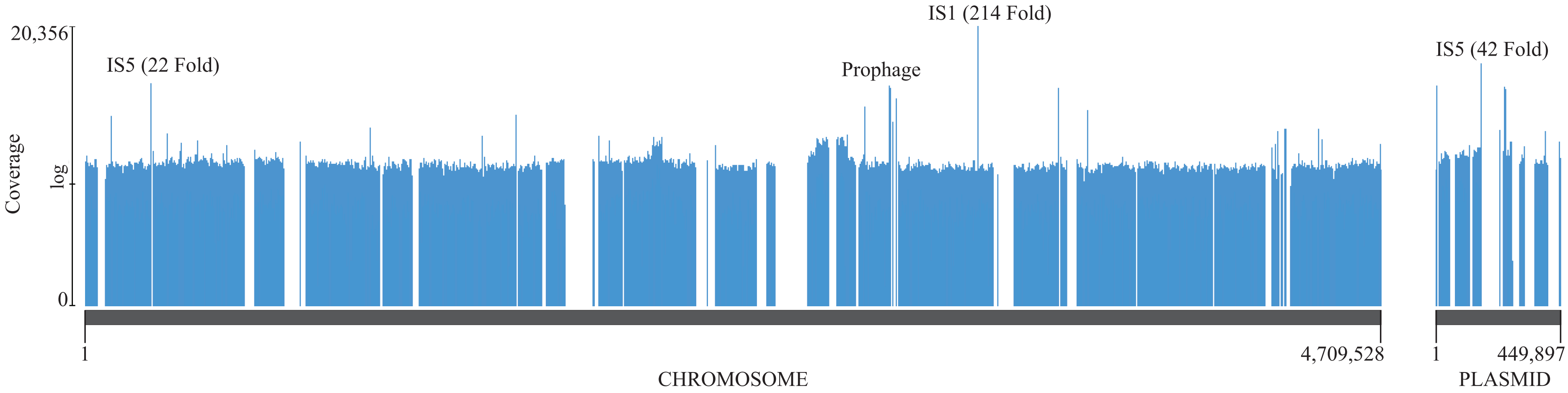


**Fig. S3.** Histogram depicting alignment of Illumina reads from strain HZ$^T$ to the *Sodalis praecaptivus* strain HS$^T$ chromosome sequence (GenBank accession CP006569), and plasmid (GenBank accession CP006570). Read coverage is depicted on a log scale, using the "log scale" setting in Geneious, highlighting overrepresentation of mobile DNA elements (prophage and IS element sequences, identified by BLAST analysis) that have undergone proliferation in the strain HZ$^T$ genome. Estimates of fold level proliferation are depicted for the most abundant elements (e.g., IS1 at 214 fold). The alignment shows that many deletions have taken place in the strain HZ$^T$ genome, consistent with functional degeneration in the transition to symbiotic life. Regarding the chromosome, strain HZ$^T$ shares approximately 4.72 Mb of DNA with *S. praecaptivus* strain HS$^T$. Regarding the plasmid, strain HZ$^T$ shares approximately 0.45 Mb of DNA with *S. praecaptivus* strain HS$^T$.

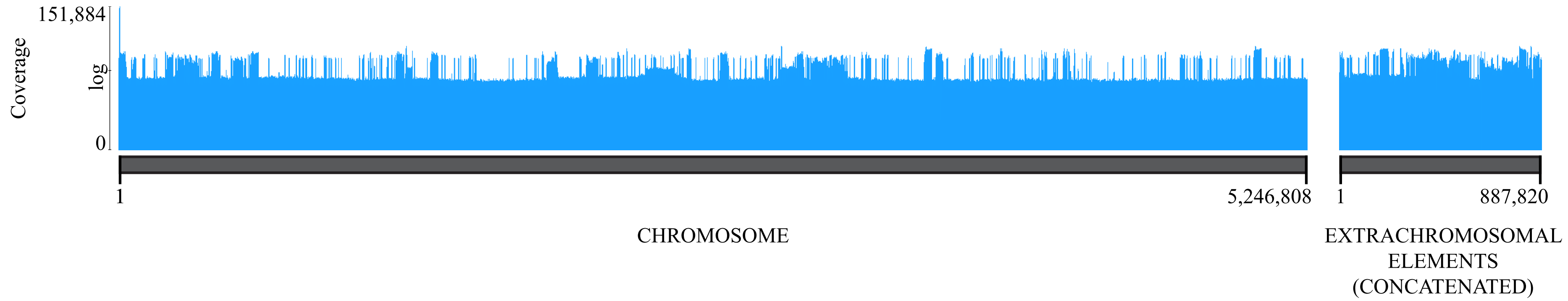


**Fig. S4.** Histogram depicting mapping of Illumina reads from strain HZ$^T$ to the final (error corrected) Nanopore assembly of the strain HZ$^T$ genome. Reads were mapped using MiniMap2 (v2.24) to a concatenated sequence of the strain HZ$^T$ Nanopore contigs comprising the chromosome and extrachromosomal elements. The alignment shows that the Illumina reads provide thorough coverage of the Nanopore contigs, validating their utility in error correction and for alignment to the *S. praecaptivus* strain HS$^T$ genome (Fig. S3) to facilitate assessment of shared gene content.

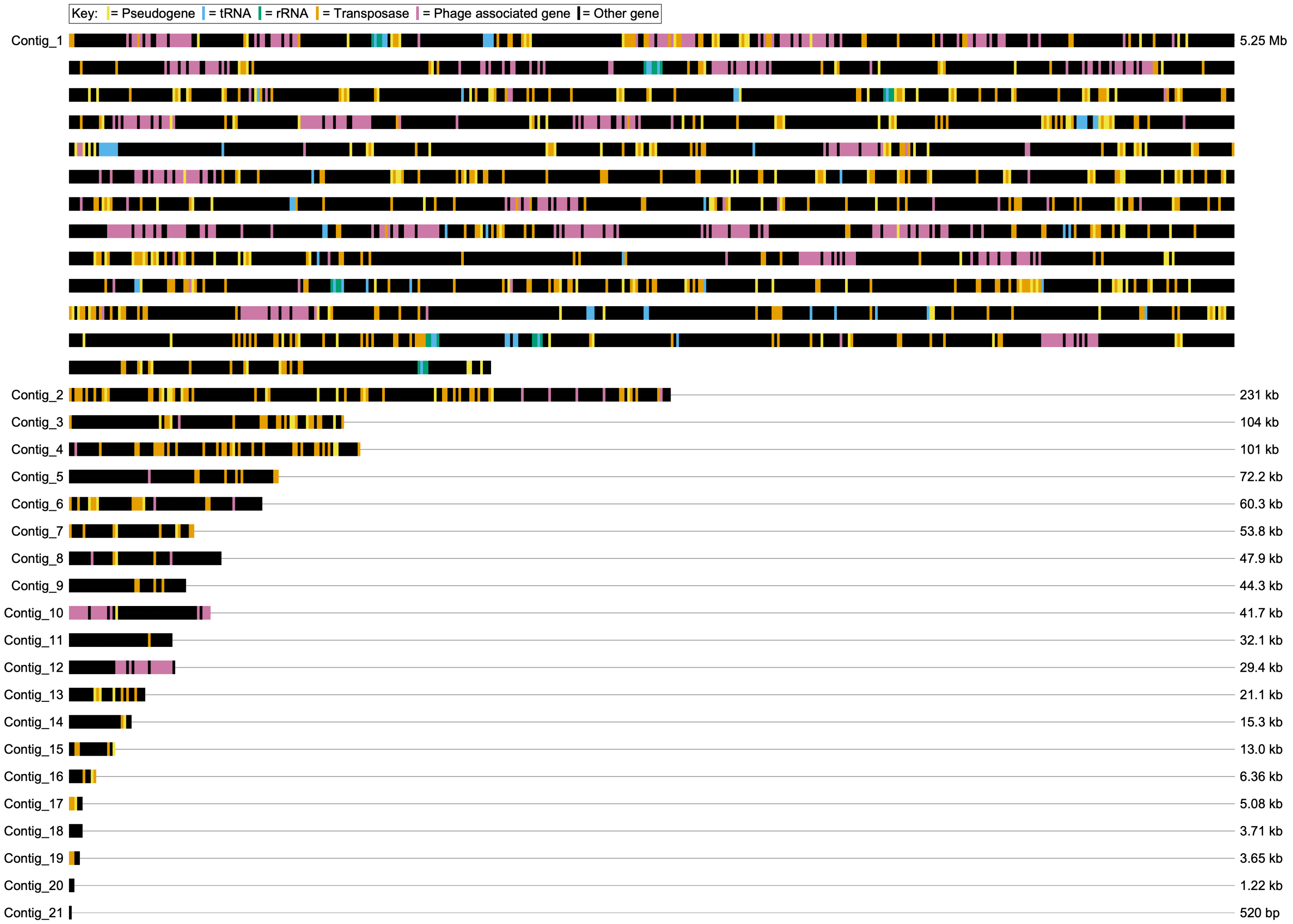


**Fig. S5.** Categorization of gene functions based on the PGAP annotation of the genome of strain HZ$^{T}$ (accession number JBDHPY000000000, and assembly number GCA_039646275.1) Genes are rendered using the same number of pixels regardless of actual size. The sizes of the contigs are presented to the right of each contig. Pseudogenes, tRNAs and rRNAs are rendered in accordance with keywords in the PGAP annotation. The keywords used to detect transposons and insertion sequences are "transposase", "tnpb", "istb", and "ista". The keywords used to detect phage genes are "phage", "tail","head", "holin", "lytic", "host specificity", "lysis", "terminase","capsid", "lysozyme", "virion", "baseplate", "rz1 lipoprotein".

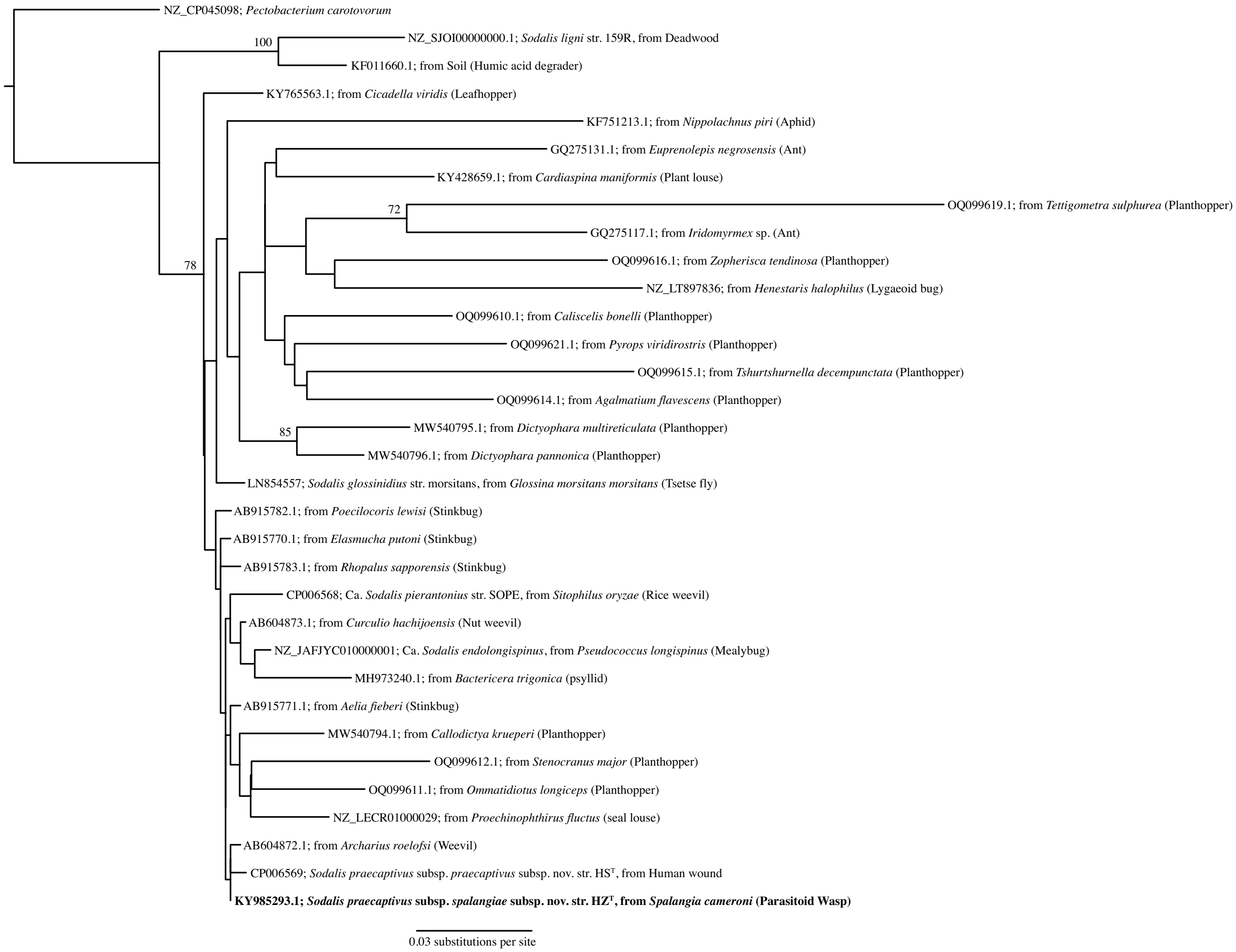


**Fig. S6.** Phylogenetic analysis of strain HZ$^{T}$, other *Sodalis* spp. and the outgroup *Pectobacterium carotovorum*. ML tree of 16S rRNA gene sequences comprising 32 *Sodalis* spp. and *Sodalis*-allied strains, including strain HZ$^{T}$ (bold), *S. praecaptivus* strain HS$^{T}$ and non insect-associated representatives from soil and deadwood.

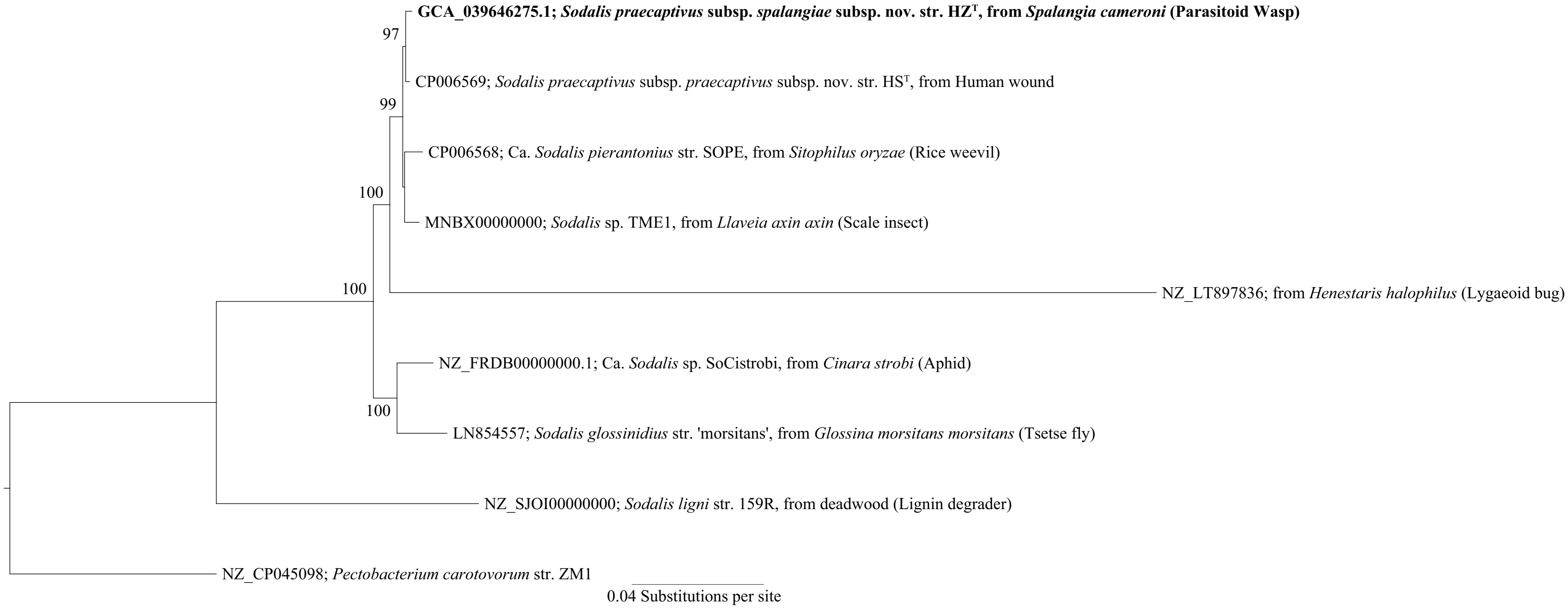


**Fig. S7.** ML tree of a subset of *Sodalis* spp. and *Sodalis*-allied strains with sequenced genomes including strain HZ$^{T}$ (bold), based on 47 conserved genes encoding ribosomal proteins, using *Pectobacterium carotovorum* as outgroup. Bootstrap support is shown only for nodes with support greater than 70%.

| | *Sodalis praecaptivus* subsp. *praecaptivus* subsp. nov. | *Ca. Sodalis pierantonius* | *Sodalis glossinidius* | *Sodalis* sp. TME1 | *Ca. Sodalis* SoCistrobi | NZ_LT897836 | *Sodalis ligni* | *Pectobacterioum carotovorum* |
|---|---|---|---|---|---|---|---|---|
| ***Sodalis praecaptivus* subsp. *spalangiae* subsp. nov.** | **0.003** | 0.009 | 0.034 | 0.008 | 0.030 | 0.238 | 0.138 | 0.184 |
| *Pectobacterioum carotovorum* | 0.183 | 0.187 | 0.195 | 0.186 | 0.190 | 0.409 | 0.204 | |
| *Sodalis ligni* | 0.137 | 0.141 | 0.149 | 0.140 | 0.145 | 0.363 | | |
| NZ_LT897836 | 0.238 | 0.241 | 0.259 | 0.240 | 0.255 | | | |
| *Ca. Sodalis* SoCistrobi | 0.028 | 0.033 | 0.026 | 0.032 | | | | |
| *Sodalis* sp. TME1 | 0.007 | 0.010 | 0.036 | | | | | |
| *Sodalis glossinidius* | 0.033 | 0.037 | | | | | | |
| *Ca. Sodalis pierantonius* | 0.008 | | | | | | | |

**Fig. S8.** Pairwise matrix of patristic distances derived from the protein-coding supergene phylogeny (Fig. S6).
